# ReporterSeq reveals genome-wide determinants of proteasome expression

**DOI:** 10.1101/2021.08.19.456712

**Authors:** Jeremy J. Work, Brian D. Alford, Annisa Dea, Asif Ali, David Pincus, Onn Brandman

## Abstract

The ubiquitin-proteasome system (UPS) is critical for cellular and organismal health. To uncover mechanisms regulating the UPS in normal and stress conditions, we systematically probed the genome of the eukaryotic model system *Saccharomyces cerevisiae* for modulators of the UPS master regulator Rpn4 under basal and stress conditions using the reverse genetic method ReporterSeq. The top UPS regulators were the thioredoxin reductase Trr1 and proteins of the large ribosomal subunit, both of which had no previously known role in UPS regulation. Unlike all known mechanisms for Rpn4 regulation which regulate Rpn4 levels, we found that Trr1 modulates the molecular activity of Rpn4 and does so in response to oxidative stress. Our work illuminates the genetic landscape through which cells regulate the UPS, and provides insight into how cells combat proteotoxicity.

## Introduction

Detection and degradation of defective proteins by the ubiquitin-proteasome system (UPS) is critical for the maintenance of a functional and healthy proteome (Leeker, Goldberg, and Mitch 2006). In order to match UPS capacity with cellular requirements, eukaryotic cells have evolved stress-activated transcriptional regulation of UPS-related genes, termed the proteolytic stress response (PSR) (Hamazaki and Murata 2020). UPS dysregulation has been associated with neurodegenerative diseases including Huntington’s disease, Alzheimer’s disease, and Parkinson’s disease, as well as cancer and aging (Q. Zheng et al. 2016). Understanding regulation of the PSR in normal and disease contexts may thus yield insights into the cause of and treatments for disease.

In budding yeast, the PSR is driven by the transcription factor Rpn4, which recognizes a specific nucleotide sequence called the proteasome-associated control element (PACE) that is found in the promoters of proteasome components, ubiquitin ligases, and other UPS factors (Mannhaupt et al. 1999; Shirozu, Yashiroda, and Murata 2015). Rpn4 continuously undergoes proteasome-mediated degradation via a ubiquitin-independent degron sequence at its N-terminus and a ubiquitylation signal near the middle of its primary sequence that is targeted by the ubiquitin ligase Ubr2 (Ju and Xie 2004; Ju et al. 2007). Competition between UPS substrates, including Rpn4, for degradation thus creates a negative feedback loop that adjusts UPS levels according to substrate load. Furthermore, several stress responses regulate *RPN4* mRNA levels, including the heat shock response (HSR, driven by Hsf1), the oxidative stress response (OSR, driven by Yap1), and the pleiotropic drug response (driven by Pdr1/3)(Ma and Liu 2010; Morimoto 2011; Lee et al. 1999; Moye-Rowley 2003).

To systematically uncover regulators of Rpn4, we used a recently developed reverse genetic approach called ReporterSeq. ReporterSeq uses a pooled assay to measure the expression of a reporter gene under CRISPR knockdown of each gene in the genome (Alford et al., 2021). Here we used ReporterSeq to identify regulators of a reporter gene targeted by Rpn4, thus providing insight into PSR regulation. The top regulators we identified were Trr1 and proteins of the large ribosomal subunit. The thioredoxin pathway regulates the PSR through a novel mechanism that targets Rpn4 molecular activity rather than its transcription or degradation. Additionally, we hypothesize that the primary substrates of the proteasome are orphan ribosomal proteins. Together, our findings reveal the genetic landscape by which cells respond to proteotoxicity and activate the PSR.

## Results

### ReporterSeq uncovers activators and suppressors of the PSR in basal conditions

To probe the genome for regulators of the PSR, we used ReporterSeq, a reverse-genetic method we recently developed to measure the effect of knocking down each gene in the genome on the activity of a specific transcription factor, in this case Rpn4 (Alford et al. 2021). We implemented ReporterSeq using a previously designed reporter for Rpn4 activity (Work and Brandman 2021) consisting of four PACE sequences followed by a crippled *CYC1* promoter, combined with a small guide RNA (sgRNA) locus expressed from the *SNR52* promoter (Figure 1a). A full list of the sgRNA sequences used can be found in Supplementary Data File 1. We created a pooled library of matched perturbation/reporter constructs spanning the genome and transfected this library into yeast expressing catalytically deactivated Cas9 (dCas9)(Gilbert et al. 2013). The pool of sgRNAs then guide dCas9 to a target locus, blocking transcription via CRISPR interference (CRISPRi). We then collected total mRNA and plasmid DNA and performed high-throughput sequencing to identify enriched and depleted reporter barcodes in two biological replicates. Finally, we computed average neighborhood normalized scores (NNS’s) of the two replicates that reflect the effect of a given gene knockdown on reporter mRNA levels (Supp. Data File 2, cols 1-2; Supp. Figure 1)(Alford et al. 2021). Genes with positive NNS values are considered PSR suppressors (their knockdown increases the PSR), while genes with negative values are identified as activators (their knockdown reduces the PSR). Mean fold change values of reporter RNA abundance for the genes with the 20 highest and lowest NNS’s can be found in Supplementary Data File 3. NNS values for individual sgRNAs can be found in Supplementary Data File 4.

**Figure 1:**
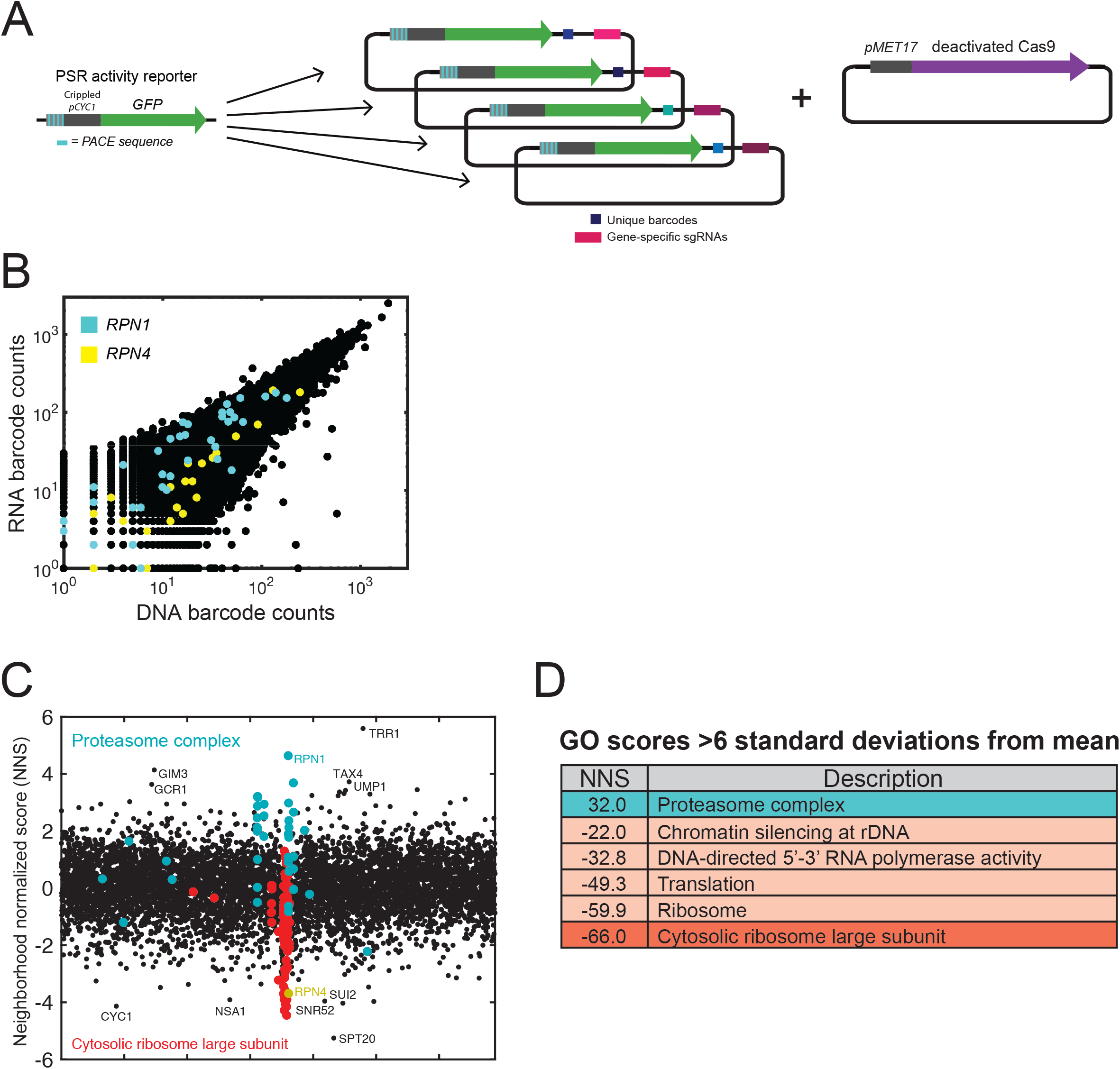
ReporterSeq uncovers activators and suppressors of the PSR in basal conditions. A: Scheme of the ReporterSeq pooled screening technology. A PSR activity reporter is cloned into a library of uniquely barcoded sgRNAs, which is coexpressed with a deactivated Cas9 construct. B: Sequencing barcode counts for RNA and DNA samples collected from the pooled library in one biological replicate. *RPN1* (cyan) and *RPN4* (yellow) are controls for increased and decreased PSR, respectively. C: Plot of neighborhood-normalized scores (NNS’s) for all gene knock-downs, arranged by alphabet. The magnitude of the NNS represents the confidence of that gene causing an effect on PSR activation. D: List of all GO categories more than six standard deviations from the mean. “Proteasome complex” and “Cytosolic ribosome large subunit” are highlighted in panel C.

We predicted that knocking down known PSR suppressors would result in enrichment of PSR reporter mRNA, while knocking down PSR activators would result in depletion. In agreement with this, expression of sgRNAs targeting *RPN1*, a subunit of the proteasome that degrades Rpn4 and is therefore an Rpn4 suppressor, increased levels of PSR reporter mRNA relative to DNA in the pooled sample (NNS=4.62, Figure 1b). Conversely, we observed that knockdown of *RPN4* resulted in decreased PSR reporter mRNA as expected (NNS=-3.71).

Among genes with the highest NNS’s were known components of the UPS (*RPN1*=4.62, *RPT1*=3.67, *UMP1*=3.29, *PRE8*=3.19, etc., Figure 1c). However, many high scoring genes, including the top overall scoring gene, *TRR1* (NNS=5.59), have no known role in the UPS (e.g. G/M3=4.14, a prefoldin subunit; *TAX4*=3.72, a protein that regulates phosphatidylinositol 4,5-bisphosphate levels and autophagy; and *GCR1*=3.63, a transcriptional activator of glycolysis genes). Other knockdowns of thioredoxin-related genes besides *TRR1* also showed positive scores for PSR activation *(AHP1*=1.71, *TSA1*=1.61, *TSA2*=0.85), suggesting that this pathway regulates the PSR. Among the lowest scores were two gene targets that matched the PSR reporter sequence (*CYC1*=-4.13, *SNR52*=-3.96), *RPN4* (−3.71), and, unexpectedly, a preponderance of genes that encode a component of the ribosomal large subunit (*RPL37A, RPL 18A, RPL36B, RPL27A*, etc.). To systematically identify the cellular pathways that contribute to PSR regulation, we computed NNS-based enrichment scores for gene ontology (GO) categories that use a Student’s t-test to compare the NNS values of their constituent genes to the set of all genes (Figure 1d)(Ashburner et al. 2000). In agreement with our observations of individual genes, the highest scoring category was “proteasome complex,” while the lowest scoring category was “cytosolic large ribosomal subunit.” We conclude these categories of genes are the most prominent suppressors (proteasome) and activators (large ribosomal subunit) of the PSR.

### Stress-specific regulators of the PSR

We performed ReporterSeq on the PSR under proteotoxic stress conditions in order to identify stress-specific regulators of the PSR. To identify these regulators, we compared the basal NNS values of all gene knockdowns to their NNS’s under proteotoxic stressors (Supp. Figure 2; Supp. Data File 1, cols 3-6). We treated our pooled knockdown library with bortezomib, a proteasome inhibitor and robust activator of the PSR (Figure 2a, left)(Work and Brandman 2021). Gene perturbations that basally activated the PSR generally blocked additional activation under bortezomib (coef=-0.37). *PDR5*, encoding a drug efflux pump, was a prominent exception, as its knockdown did not affect basal PSR activity, yet it still blocked PSR activation under bortezomib. This observation is in line with previous work showing that drug efflux pumps reduce the efficacy of bortezomib treatment (Fleming et al. 2002). We next treated the knockdown library with hydrogen peroxide, an oxidizing agent, and found that the interactions were also anti-correlated with the basal NNS scores (coef=-0.51, Figure 2a right), suggesting that PSR activity buffers against oxidative insults. An exception to this was knockdown of *TRR1*, which had a higher interaction score than expected based on its basal score. *TRR1* disruption causes oxidative stress (Carmel-Harel et al. 2001), which is likely exacerbated by hydrogen peroxide, and suggests that PSR activation is an important response to oxidative stress. Conversely, knockdown of *YAP1* blocked PSR activation under hydrogen peroxide treatment, matching previous characterization of Yap1-mediated Rpn4 regulation (Lee et al. 1999). NNS values for individual sgRNAs under each stressor can be found in Supplementary Data File 5.

**Figure 2:**
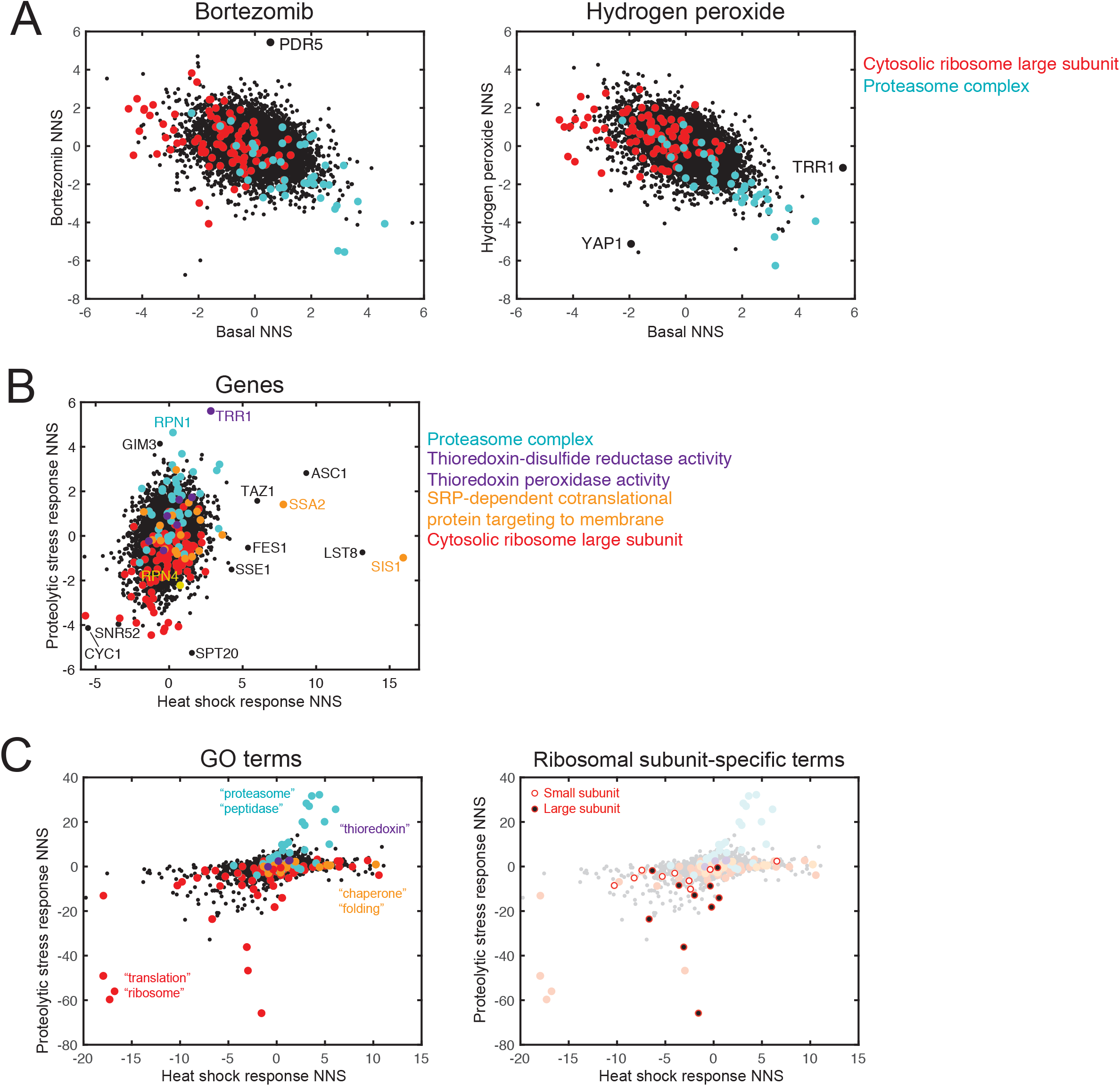
The PSR and HSR are regulated by non-overlapping pathways. A: Plot of all gene NNS’s in basal conditions vs. 3 hr 20 μM bortezomib (left) or 30 min 200 μM hydrogen peroxide treatment (right). Two GO categories are highlighted with color. B: Plot of all gene NNS’s for both the PSR and HSR, with *RPN4* and several GO categories highlighted with color. C: Plot of all GO term NNS’s for both the PSR and HSR, with groups containing the indicated words highlighted.

### The PSR and HSR are regulated by non-overlapping pathways

Previous work has used ReporterSeq to identify regulators of the HSR (Alford et al. 2021). Both the HSR and the PSR are central protective mechanisms against proteotoxic stress, and they are commonly thought to work in parallel to repair or eliminate misfolded proteins, respectively. With this in mind, we investigated whether the HSR and PSR have common activators and repressors, or if they are instead activated by unique sets of factors and pathways. We compared our collection of PSR ReporterSeq gene scores to the previously published data set for the HSR that used the same library of sgRNAs (Figure 2b). The two responses were weakly correlated (coef=0.22); the two most prominent regulating GO categories of the PSR (“proteasome complex” and “cytosolic large ribosomal subunit”) had only minor effects on the HSR, while a prominent HSR-regulating group (“chaperone cofactor-dependent protein refolding”) minimally affected the PSR. Consistent with this, the top regulator of the HSR (*SIS1*, NNS_HSR_= 16.0) negatively affected the PSR (NNS_PSR_= -1.00), although the top regulator of the PSR *(TRR1*, NNS_PSR_= 5.59) regulated the HSR to a lesser extent (NNS_HSR_=2.86). To further compare HSR and PSR regulation, we plotted the enrichment scores of GO categories (Figure 2c). Both responses were reduced by knockdown of ribosome-related genes, but these were specific to different subunits: the large subunit in the case of the PSR, and the small ribosomal subunit for the HSR. The vast majority of top PSR-regulating groups contained “proteasome” or “peptidase” in their descriptors, and scored much higher for PSR activation than HSR activation. Conversely, groups containing “chaperone” or “folding” in their descriptors were among the highest HSR-activating groups, but all scored near zero for the PSR. These observations highlight a lack of overlap between genetic regulators of the HSR and PSR, and support previous work finding that folding stressors minimally activate the PSR and proteolytic stressors minimally activate the HSR (Work and Brandman 2021).

### Ribosome biogenesis generates demand for the PSR

Since protein degradation depends on protein synthesis, it was expected that knockdowns of genes involved in translation and the ribosome would reduce PSR activity. However, we found specific enrichment for the genes encoding proteins in the large ribosomal subunit as well as genes involved in rDNA regulation (Figure 1d). Together, these categories suggest that the PSR may play a role in ribosome biogenesis. One way the PSR could be involved in ribosome biogenesis is by degrading ribosomal proteins that are produced in stoichiometric excess. Imbalances in the stoichiometry of ribosomal proteins trigger a specific stress response (Tye et al. 2019; Albert et al. 2019), and the UPS is known to enforce appropriate stoichiometry by degrading supernumerary ribosomal proteins (Sung, Reitsma, et al. 2016; Sung, Porras-Yakushi, et al. 2016). When expression of individual ribosomal protein genes is reduced or when unincorporated ribosomal proteins persistently accumulate, cells are known to adjust and globally downregulate ribosome biogenesis (Zhao, Sohn, and Warner 2003; Reiter et al. 2011; Albert et al. 2019). Thus, we hypothesized that cells in which ribosomal protein expression is reduced by CRISPRi may produce fewer excess ribosomal proteins and thereby have reduced demand for the UPS.

To test whether reduced expression of ribosomal proteins decreases PSR activity, we took advantage of the drug rapamycin. Rapamycin inhibits the Target of Rapamycin Complex I (TORC1) kinase, and yeast lack the TORC1/4E-BP translational control pathway present in mammalian cells. The primary function of TORC1 in yeast is activation of ribosome biogenesis at the transcriptional level (Powers and Walter 1999; Crespo and Hall 2002; Martin, Soulard, and Hall 2004). Thus, rapamycin specifically results in the downregulation of ribosomal protein genes and other ribosome biogenesis factors while initially having few effects on the remaining transcriptome or translation (Reiter et al. 2011; Zencir et al. 2020). To assess the role of ribosome biogenesis in PSR activation, we treated wild type cells expressing the PSR reporter with bortezomib in the presence and absence of rapamycin and performed a time course to examine both the immediate and sustained effects of rapamycin. Following bortezomib treatment, cells began to induce the PSR after 1 hour and reached a plateau after 3 hours (Figure 3a). Pretreatment with rapamycin for 30 minutes modestly increased basal PSR activity, but blunted the response to bortezomib to a shallow plateau, consistent with lower Rpn4 levels and excess proteasome capacity. To more directly test the capacity of the proteasome, we treated cells expressing two fluorescent degron reporters, ERm-Deg, and Cyto-Deg, which consist of a superfolder GFP fused to a 23-33 amino acid degron sequence (Work and Brandman 2021; Maurer et al. 2016). These reporters are expression-controlled by an upstream mCherry that is connected to the GFP reading frame by two tandem T2A sequences, which cause the mCherry to detach from the ribosome before the GFP begins translation (Donnelly et al. 2001; Szymczak and Vignali 2005). Rapamycin reduced the stability of both degron reporters (Figure 3b, left). In the presence of rapamycin, the highest dose of bortezomib tested stabilized the degrons to just their untreated levels (Figure 3b, right). With the caveat that sustained TORC1 inhibition results in reduced ribosomes, these data suggest that blocking ribosome biogenesis reduces demand for the PSR, and we propose that ribosomal proteins are a major category of proteasome clients.

**Figure 3:**
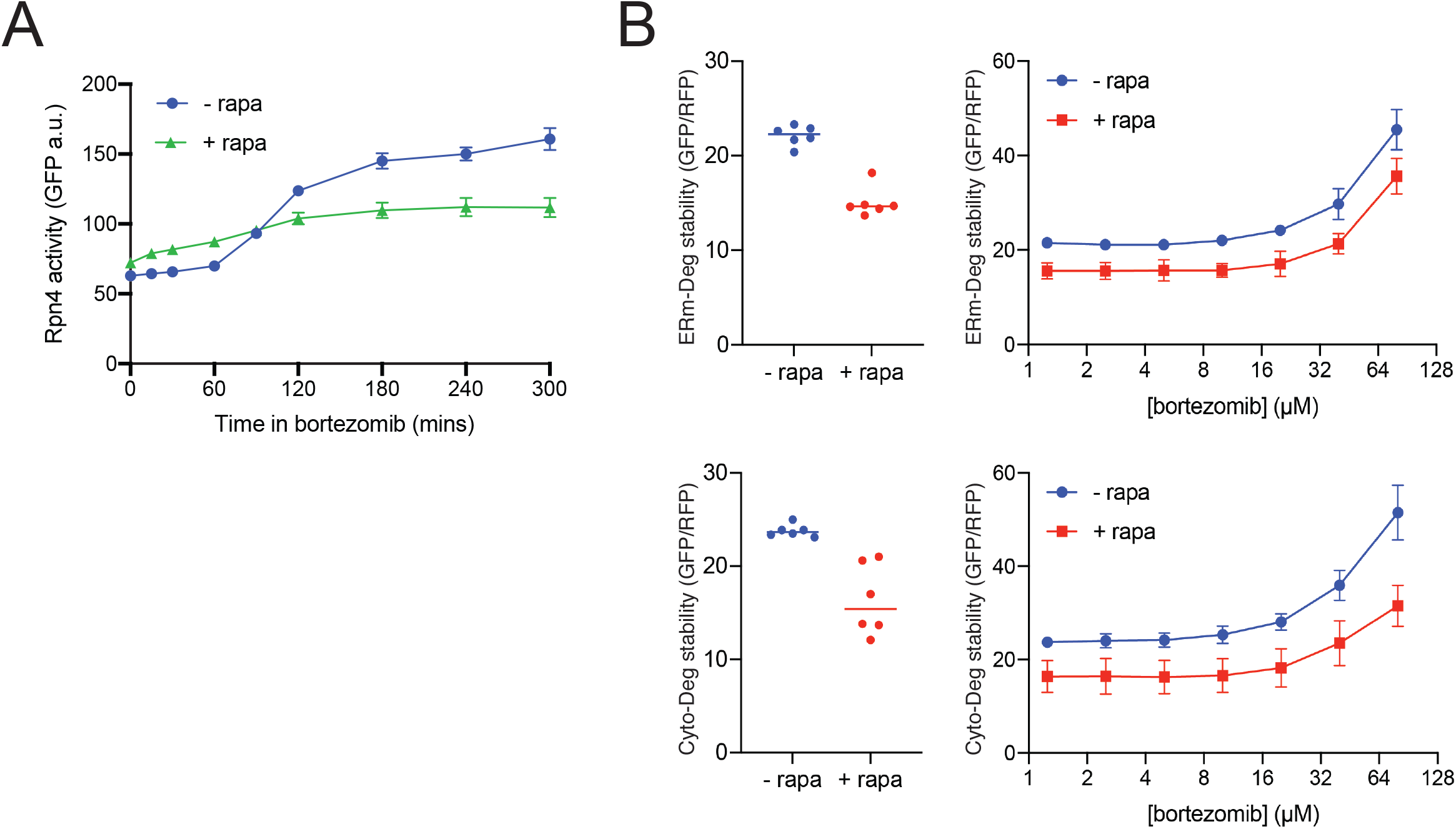
Ribosome components burden the proteasome. A: Plot of Rpn4 activity over time under 40 μM bortezomib with or without 30 min pretreatment with 10 μM rapamycin. B: Plot of ERm-Deg (top) and Cyto-Deg (bottom) stability under basal conditions (left) or a titration of bortezomib (right), with or without 30 min pretreatment with 10 μM rapamycin. These reporters are structured as RFP-(T2A)_2_-GFP-degron, which allows RFP to serve as a same-cell expression control for the GFP-degron species. rapa, rapamycin.

### Trr1 regulates Rpn4 molecular activity

As the top regulator of the PSR with no known role in UPS regulation, we investigated the role of *TRR1* in suppression of Rpn4. To lower expression of *TRR1* in followup assays, we created a genomically integrated TRR1-knockdown strain, in which three CGA codons were added to the 3’ end of the *TRR1* coding sequence prior to the stop codon (Supp. Figure 4a). Repeats of the CGA codon result in stalled translation and recruitment of the ribosome quality control complex when present in tandem in an **mRNA** coding sequence, reducing the full-length levels of a tagged protein by 93% (Letzring, Dean, and Grayhack 2010). Consistent with the screen results, our TRR1-knockdown strain exhibited a 3-fold activation of Rpn4 (Figure 4a). This level of activation was similar to that of treatment with 10 µM bortezomib, deletion of *UMP1* (a proteasome assembly chaperone), or deletion of *UBR2*.

**Figure 4:**
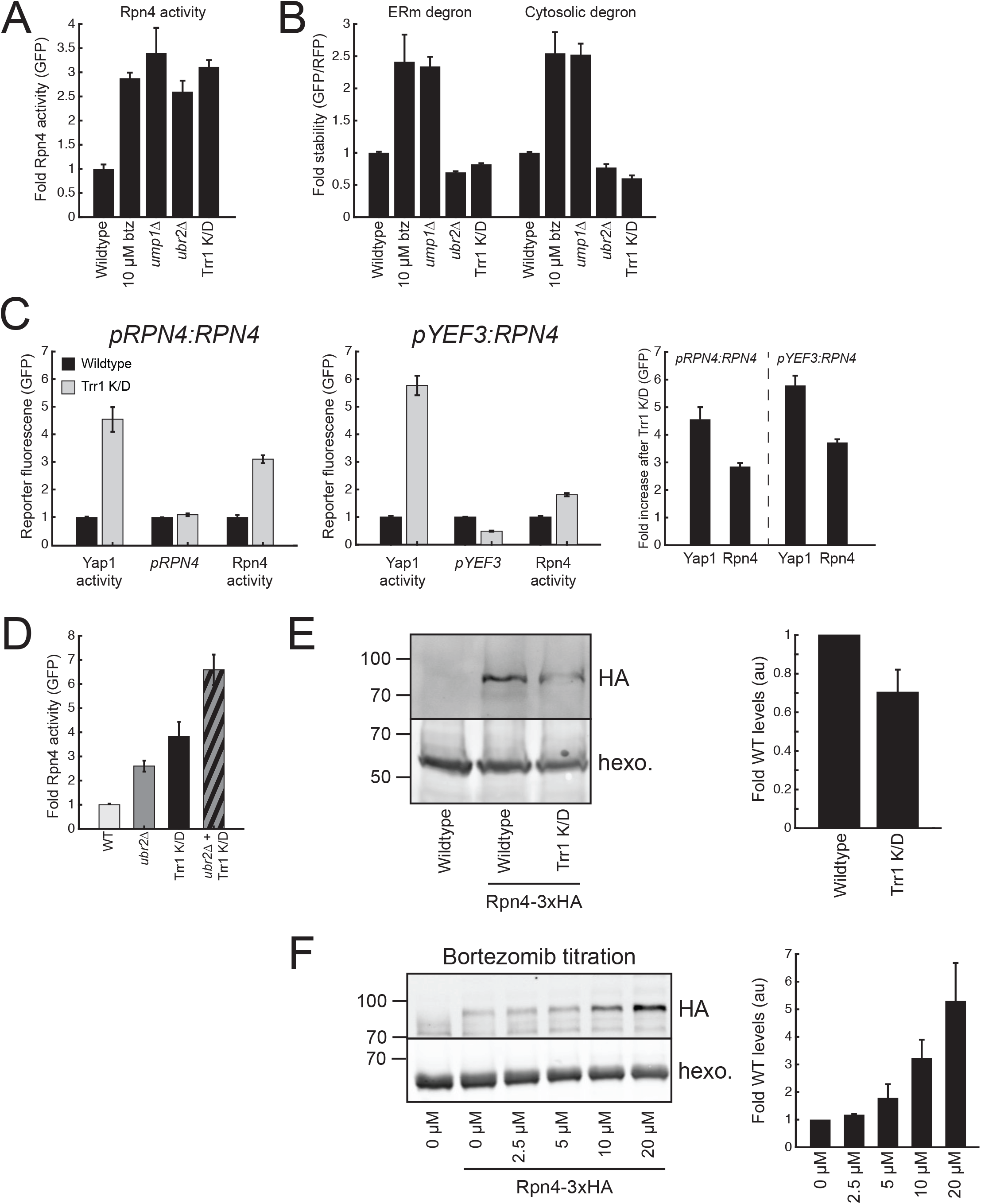
Trr1 regulates the molecular activity of Rpn4. A: Measurement of Rpn4 activity upon Trr1 knockdown, normalized to wildtype. B: Measurement of the stability of ERm-Deg and Cyto-Deg, two differently-localized degron reporters. C: Measurement of Yap1, *pRPN4*, and Rpn4 activity upon Trr1 knockdown in either a wildtype *RPN4* (left) or *pYEF3:RPN4* (center) background. These values are combined to provide the fold-increase after Trr1 knockdown (right). D: Rpn4 activity upon Ubr2 deletion, Trr1 knockdown, or both. E: (left) Representative immunoblot of HA-tagged Rpn4 upon Trr1 knockdown, with quantification of n=3 measurements (right). F: (left) Representative immunoblot of HA-tagged Rpn4 upon bortezomib treatment, with quantification of n=3 measurements (right). All error bars denote standard error of measurement, n≥3.

Two broad hypotheses may explain how low Trr1 levels activate Rpn4: 1) low Trr1 levels result in widespread protein damage and new UPS substrates that outcompete Rpn4 for degradation, leading to increased Rpn4 levels and resulting PSR activity, and 2) Trr1 more directly regulates Rpn4 activity via its synthesis, degradation, or molecular activity. To distinguish between these hypotheses, we measured the stability of ERm-Deg and Cyto-Deg upon Trr1 knockdown. We predicted that under hypothesis #1, the stability of these reporters would increase due to competition with other substrates, a state that can be recapitulated with bortezomib or *ump*Δ. Conversely, under hypothesis #2, degron stability would decrease due to increased proteasome expression via Rpn4 in the absence of stress, similarly to deletion of *UBR2*, another Rpn4 regulator. Both ERm-Deg and Cyto-Deg decreased in stability upon knockdown of Trr1 (Figure 4b), and we therefore conclude that Trr1 regulates Rpn4 synthesis, degradation, and/or molecular activity without increasing UPS substrates.

Previous work studying Rpn4 has shown that Trr1 undergoes transcription-level regulation by upstream stress-dependent transcription factors such as Hsf1, Yap1, and Pdr1/3 (Hahn, Neef, and Thiele 2006; Temple, Perrone, and Dawes 2005; Moye-Rowley 2003). Based on this work, a plausible mechanism for Trr1-mediated regulation of Rpn4 is via Yap1 activation. Indeed, Yap1 is known to respond to thioredoxin perturbation and oxidative stress, and it targets both *TRR1* and *RPN4* for transcription as part of the OSR (lzawa et al. 1999; Delaunay, lsnard, and Toledano 2000; Lee et al. 1999). To test this hypothesis, we measured the effect of *TRR1* knockdown on reporters for OSR activity and *RPN4* transcription. We constructed the OSR activity reporter similarly to the PSR activity reporter except the Yap1 binding sequence “TTACTAA” was encoded upstream of the crippled *CYC1* promoter fragment rather than the Rpn4 binding sequence (Supp. Figure 4b)(Fernandes, Rodrigues-Pousada, and Struhl 1997). *RPN4* transcription was measured using a previously designed reporter featuring GFP expressed from the *RPN4* promoter (Work and Brandman 2021). Upon Trr1 knockdown, Yap1 and Rpn4 activity increased to 4.5-fold and 3-fold, respectively; however, *RPN4* transcription only activated to 1.1-fold of basal levels (Figure 4c left). Replacing the native *RPN4* promoter with the constitutive *YEF3* promoter still permitted full induction of Rpn4 upon Trr1 knockdown, despite lacking any Yap1 binding sites (Figure 4c). We conclude that Trr1-mediated regulation of Rpn4 does not occur through transcriptional targeting of *RPN4*.

Rpn4 also undergoes degradation-level regulation via a ubiquitylation site recognized by the E3 ligase Ubr2 (Ju et al. 2007) and a ubiquitin-independent degron signal at the N-terminus (Ju and Xie 2004). A previous study observed that removing the ubiquitin-independent signal minimally affects Rpn4 half-life (Ju and Xie 2004), so we hypothesized that if Trr1-mediated regulation of Rpn4 functions through degradation, it likely does so through regulation of the Ubr2-mediated pathway. We tested for genetic interactions between Ubr2 deletion and Trr1 knockdown and found that while both perturbations increase Rpn4 activity, their combined perturbation was additive, and thus there was no significant interaction measured by a two-way ANOVA test (p=0.24, Figure 4d). We conclude that Trr1 does not regulate Ubr2-mediated Rpn4 degradation.

While previous studies of Rpn4 have identified mechanisms that regulate Rpn4 protein levels, whether the molecular activity Rpn4 is modulated has not been investigated. To explore this, we measured the protein levels of HA-tagged Rpn4 in tandem with PSR activity. Surprisingly, while Trr1 knockdown caused an increase in Rpn4 activity, it caused a decrease in Rpn4 protein levels (Figure 4e). This contrasts with bortezomib-induced Rpn4 activity, which can be characterized by an increase in Rpn4 levels in a dose dependent manner (Figure 4f). We conclude that Trr1 knockdown causes increased Rpn4 molecular activity.

### The thioredoxin system activates Rpn4 in response to oxidative stress

We next investigated whether regulation of Rpn4 molecular activity is specific to Trr1 knockdown or is part of a larger pathway. Deletion of Tsa1, a thioredoxin peroxidase whose knockdown also led to PSR activation in our ReporterSeq screen (NNS=1.61), caused a ∼2-fold increase in Rpn4 activity compared to wildtype, with no significant change to levels (Figure 5ab). Therefore, disruption of other components of the thioredoxin system are sufficient to increase Rpn4 molecular activity, presumably through the same mechanism employed upon Trr1 knockdown. We next tested whether perturbation of the glutathione system, a redox system that operates in parallel to the thioredoxin system (Grant 2001), is sufficient to activate Rpn4. We knocked down the glutathione regulators Gsh1, Gpx1, or Gpx2 through the addition of three CGA codons and found that this led to *reduced* Rpn4 activity rather than the increased activity observed in thioredoxin hypomorph strains (Figure 5c). Thus, the thioredoxin system is unique in regulating Rpn4. Finally, we tested whether general oxidative stress activates Rpn4. We treated cells with 200 µM hydrogen peroxide, a known activator of Yap1 (Fernandes, Rodrigues-Pousada, and Struhl 1997), and found that Rpn4 was robustly activated to 2.5-fold, which was comparable to the activation exhibited upon treatment with 10 µM bortezomib (Figure 5a). However, we observed that Rpn4 levels are not significantly changed upon peroxide treatment (Figure 5b), even though they increased 3-fold under bortezomib treatment. Indeed, when comparing the activity and levels of Rpn4 upon Trr1 knockdown, Tsa1 deletion, or peroxide treatment to a bortezomib standard curve, we observed that all three redox perturbations resulted in a higher Rpn4 molecular activity than that of bortezomib treatment. Furthermore, we found that deletion of Yap1 did not impair Rpn4 activation under hydrogen peroxide (Figure 5d). Hydrogen peroxide also decreased the stability of ERm-Deg and Cyto-Deg, suggesting that it does not impair the proteasome or cause proteolytic stress (Figure 5e). We conclude that the thioredoxin system and oxidative stressors control the PSR via the molecular activity of Rpn4, independently from Yap1 or proteasomal degradation (Figure 6).

**Figure 5:**
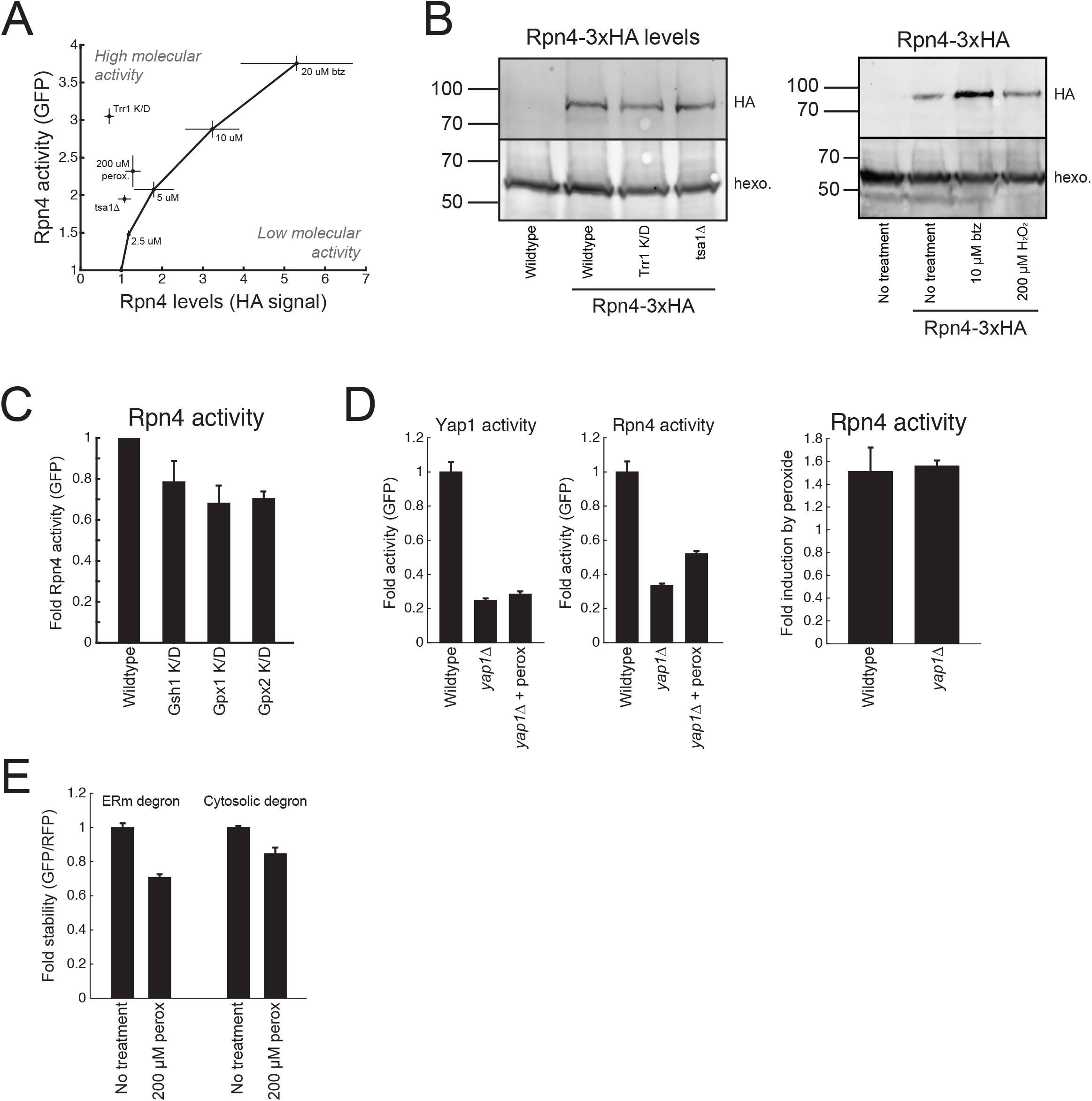
The thioredoxin system activates Rpn4 in response to oxidative stress. A: Plot of Rpn4 activity vs. Rpn4 levels for several perturbations, including a titration of bortezomib. B: Representative immunoblot of HA-tagged Rpn4 upon Trr1 knockdown or Tsa1 deletion (left) or treatment with bortezomib or hydrogen peroxide (right), with quantification of n=3 measurements in (A). C: Rpn4 activity upon knockdown of glutathione system factors, Gsh1, Gpx1, and Gpx2. D: Yap1 (left) and Rpn4 (middle) activity in the noted backgrounds and drug treatments, fold Rpn4 activity induction upon treatment with 200 μM hydrogen peroxide, normalized to no treatment in the noted background in a same day experiment (right). Hydrogen peroxide treatment: 200 μM. All treatments were for 5 hrs. All error bars denote standard error of measurement, n≥3. btz, bortezomib; perox, hydrogen peroxide.

**Figure 6:**
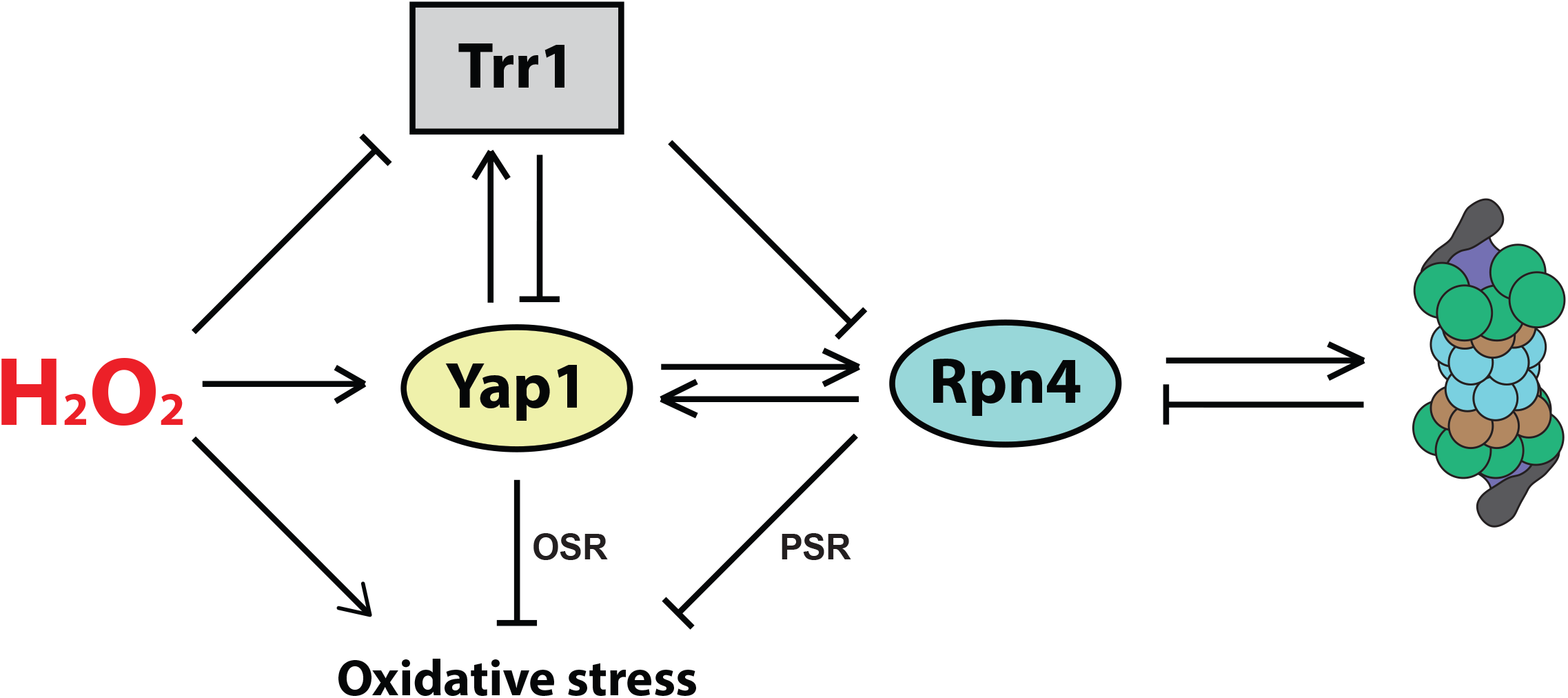
Model for Trr1-mediated activation of Rpn4 in response to oxidative stress. Under this model, hydrogen peroxide causes oxidative stress, activates Yap1, and impairs Trr1. Trr1 impairment derepresses Yap1 and Rpn4, which drive expression of each other and effect the OSR and PSR, respectively, resolving the oxidative stress.

## Discussion

We used ReporterSeq, a high-throughput pooled screening technology, to identify top regulators of the PSR in yeast. Our method successfully identified several known PSR regulators, as well as many newly implicated factors, such as the ribosome large subunit and the thioredoxin system. We explored the mechanism by which the thioredoxin system regulates the PSR, and we found that contrary to the commonly attributed Yap1-Rpn4 pathway, the thioredoxin pathway directly regulates Rpn4 molecular activity. Our work provides a rich dataset that will continue to inform future work on the PSR and other related stress pathways.

The PSR and HSR are often described as tightly coupled, both because of their shared goal in resolving misfolded proteins, and because of the direct transcriptional regulation of *RPN4* by Hsf1 (Hahn, Neef, and Thiele 2006). However, our work suggests that while there are some gene knockdowns that are capable of activating both pathways (e.g. *ASC1* and *TAZ1)*, most top regulators of either pathway have minimal effects on the other. In particular, chaperone knockdowns that activated the HSR (*SIS1, SSA1, FES1, SSE1)* had minimal effects on the PSR, even though the elevated Hsf1 activity should be able to enhance *RPN4* transcription. In agreement with this, we observed in previous work that deletion of the chaperone genes *HSC82, SSA2*, and *HSP104* all activated the HSR but had no significant effect on the PSR (Work and Brandman 2021).

Not only does the PSR fail to robustly activate in folding stress, we also discovered here that Yap1-mediated transcription of *RPN4* is minimal under oxidative stress, Instead, the bulk of PSR activation in oxidative stress comes from increasing the molecular activity of Rpn4 via the thioredoxin system. Several models exist for how Rpn4 may help resolve oxidative stress. One possibility is that Rpn4 promotes the OSR through transcription of *YAP1*; not only does Yap1 target *RPN4*, but Rpn4 also targets *YAP1*, thus promoting positive feedback during stress (Ma and Liu 2010). Another possibility is that Rpn4 protects against DNA damage caused by hydrogen peroxide or thioredoxin perturbation. Oxidative stress is known to cause DNA damage, and Rpn4 targets several genes involved in DNA damage repair (Karpov et al. 2013). Intriguingly, previous work describing Yap1 targets does not identify any factors directly involved in DNA damage repair (Rodrigues-Pousada, Menezes, and Pimentel 2010; Lee et al. 1999), so Rpn4 may have been delegated this responsibility under oxidative stress. Finally, the UPS not only degrades proteins, but it has also been implicated in reprogramming the proteome through alternate ubiquitin linkages in response to oxidative stress (Silva, Finley, and Vogel 2015). This altered functionality of the UPS upon oxidation could presumably be enhanced by increased expression of UPS factors by the PSR.

How the thioredoxin system regulates the molecular activity of Rpn4 remains to be elucidated. One possibility is that Tsa1 serves as a direct inhibitor of Rpn4. Past studies have shown that Tsa1 can serve as a molecular chaperone, and this chaperoning function can be disrupted by perturbation or misregulation of Tsa1 (Hanzén et al. 2016; Trotter et al. 2008). These observations align with a model whereby Tsa1 binds and inhibits Rpn4 and would sufficiently explain why both deletion of *TSA1* and knockdown of Trr1 are able to activate Rpn4. Such inhibition would be reminiscent of the inhibition of Hsf1 by multiple chaperones (Zou et al. 1998; X. Zheng et al. 2016; Neef et al. 2014). Another possibility is that Rpn4 is posttranslationally modified by Trr1, Tsa1, or an intermediary. Rpn4 contains six cysteines, two in a C2H2 zinc finger domain (Mannhaupt et al. 1999), and four with no known functions. An intriguing possibility is that these cysteines form disulfide bonds or become otherwise modified during oxidative stress, resulting in enhanced Rpn4 molecular activity. Such a mechanism of regulation has precedent in yeast - Yap1 forms an intramolecular disulfide bond and relocalizes to the nucleus under oxidative stress (Kuge, Jones, and Nomoto 1997). Further investigation of the interplay between oxidative stressors and the PSR may advance our understanding of proteotoxicity and disease.

## Methods

### Yeast strain construction

All yeast strains were created from BY4741. Construction was done by transforming cells with a crude PCR product bearing 40 base pair overhangs homologous to the target genomic locus, a selection cassette for either antibiotic selection in YPD or auxotrophic selection in SD, and any other desired sequences. Deletion strains were constructed via transformation with antibiotic selection cassettes (NATMX6 or KANMX6), amplified with overhangs flanking the open reading frame to be deleted. Knockdown strains that made use of tandem C-terminal CGA codons were constructed via transformation with NATMX6, amplified with overhangs flanking the target gene stop codon. The forward primer was designed to include the sequence CGACGACGATAG to incorporate three knockdown codons and a new stop codon. The *pYEF3:RPN4* strain was constructed via transformation by inserting a His3 cassette and 1000nt of *p YEF3* immediately upstream of the *RPN4* open reading frame. All transformants were verified by genomic PCR.

### Reporter plasmid and sgRNA library generation

Reporter plasmids used in this study were cloned using the Gibson Assembly method and the NEBuilder HiFi DNA Assembly Master Mix (New England Biolabs). The PSR reporter, OSR reporter, *pRPN4:GFP* reporter, Cyto-Deg, and ER-Deg plasmids were expressed from high-copy 2-micron plasmids containing a Ura3 selection cassette. The ERm-Deg and Cyto-Deg sequences are “10-21” and “10-31” from Maurer et al., 2016, respectively, and are fused to GFP via a Gly-Ser-Gly-Ser linker.

The sgRNA library was created in a near-identical manner to that performed in (Alford et al. 2021), except with the PSR reporter instead of an HSR reporter. Approximately 12 sgRNAs were designed to target the 5’ end of each protein-coding and non-coding gene in the genome (between -750 and +50 nucleotides from the start of the ORF with priority toward sgRNAs closer to the ORF start) based on the Saccharomyces Genome Database (Cherry et al. 2012). These sgRNA sequences are the same as the ones used in (Alford et al. 2021). A pool of oligonucleotides was ordered (CustomArray, Inc) containing these sequences and constant flanking sequences for the purposes of cloning. A 2-micron Ura3-selective plasmid was constructed from three DNA fragments using HiFi DNA Assembly Master Mix (New England Biolabs) for cloning. The first fragment was a diverse sample of inserts amplified via PCR using Platinum Taq DNA Polymerase (Thermo Fisher Scientific), producing a sequence that contained random barcodes associated with random sgRNAs from the library. The second fragment was the parent plasmid digested at two locations using the FastDigest enzymes Bgll and Bglll (Thermo Fisher Scientific). The final fragment was another region of the plasmid amplified via PCR, which was necessary to connect the previous two pieces. These three fragments were assembled in a 100 µL reaction containing 700 ng of the first fragment, 600 ng of the second fragment, and 400 ng of the final fragment. After a 1 hr incubation at 50° C, the reaction was concentrated to 20 µL using a DNA clean and concentrate column (Zymo Research) and electroporated in five separate cuvettes containing 4 µL of the plasmid and 40 µL of ElectroTen-Blue electrocompetent cells (Agilent). After a 1 hr outgrowth in Luria Broth (LB) at 37° C, the plasmid library was then grown in 1 L of LB containing 100 µg/mL carbenicillin for 2−3 days until it reached saturation. The plasmid was then purified via maxiprep (Qiagen).

### Plasmid library sequencing

The sgRNA library sequencing was done in an identical manner to that performed in (Alford et al. 2021). Two micrograms of plasmid were digested with the FastDigest Pstl restriction enzyme (Thermo Fisher Scientific), and the 900 bp piece was gel purified using a gel DNA extraction kit (Zymo Research), then recirculized in a 20 µL reaction containing 60 ng of DNA and 1 µL of T4 DNA Ligase (Thermo Fisher Scientific) in buffer. The ligation product was then amplified with primers containing the appropriate overhangs for high-throughput sequencing and then gel purified. This piece was then sequenced through paired-end sequencing using custom primers on an lllumina MiSeq sequencing machine.

### Yeast sample preparation for screening

Yeast from a saturated library sample were grown overnight to log phase in SD -Ura/Leu. These yeast were then split into a number of cultures of at least 10 ml. Yeast were then either left untreated or stressed with 20 uM bortezomib for 3 hr, or 200 uM hydrogen peroxide for 30 min. The yeast were then vacuum filtered onto nitrocellulose paper and flash frozen in liquid nitrogen. This procedure was performed in duplicate, and each replicate came from separate transformations of the library into the yeast, separate treatments, and sample preparations. Some basal samples included technical duplicates (independent sequencing sample preparation of the same yeast sample), and these were averaged into one biological replicate.

### Nucleic acid sample collection

Collection and preparation of sample RNA and DNA was done in a near-identical manner to the protocol in (Alford et al. 2021). The yeast DNA samples were collected by thawing frozen yeast and following the standard procedure of a Zymoprep Yeast Plasmid Miniprep Kit (Zymo Research).

RNA was extracted through acid-phenol extraction. For each sample, a frozen yeast pellet was resuspended in 450 µL AE Buffer (50 mM sodium acetate pH 5.5, 10 mM EDTA), 50 µL 10% SDS, and 500 µL acid phenol:chloroform (Thermo Fisher Scientific). This mixture was heated to 65°C and shaken at 1400 rpm for 10 min, followed by 5 min on ice. The mixture was then centrifuged at 12,000 g for 15 min. The supernatant was then added to 450 µL of chloroform and centrifuged at 15,000 g for 5 min. The nucleic acids were extracted from the supernatant through standard isopropanol extraction.

NextFlex oligo-dT beads (PerkinElmer) were used to select polyadenylated mRNA from 100 µg of the extracted nucleic acids. This sample was subjected to DNAse treatment using Turbo DNase (Thermo Fisher Scientific) treatment and then purified using AMPure XP beads (Beckman Coulter). The resulting RNA was reverse transcribed using Multiscribe reverse transcriptase and a gene-specific primer (Thermo Fisher Scientific), and the resulting cDNA was purified using alkaline lysis of the RNA, followed by another round of AM Pure bead purification.

The purified cDNA and purified DNA were then amplified at the barcode region using primers with the appropriate lllumina sequencing ends and indexing barcodes to differentiate each sample. These PCR products were then gel purified, mixed proportionately with each other, and then sequenced using a lllumina next-generation sequencing machine (NextSeq).

### Scoring and statistical analysis

NNS scores were analyzed the same way as in Alford et al., 2021. In brief, basal NNS’s and interaction NNS’s were calculated from pairs of conditions. For the case of the basal NNS values, RNA levels and DNA levels were compared, whereas for the gene-stressors interaction NNS values, stressor RNA was compared to unstressed RNA. For each comparison, A and B, and each barcode, x, an NNS was computed, score(x,A,B), which captures how extreme (positive or negative) the read count is for barcode x in A vs B. These scores account for the discrepancies between counts in different conditions, taking into account the higher relative amount of noise in small counts versus larger counts, and the fact that some conditions yield higher read counts across most barcodes. These scores for each barcode were then aggregated to form a score for each gene. The gene NNS’s were scaled such that a score of 1 represents one standard deviation from the mean. To calculate the GO category scores, two groups were compared: the NNS’s of all the genes within the given category, and the set of all NNS’s. A Student’s t-test was performed to determine if the GO category set deviates from the full set, and the natural logarithm the reciprocal p-value was reported as the NNS. NNS values reported throughout the text are averages of the NNS value attained in each of the two biological replicates sequenced.

MATLAB programs (MathWorks) and the raw data to compute gene NNS’s and GO category enrichment can be found in Supplementary Data File 6.

### Fluorescent reporter assay measurements

Yeast samples were grown overnight in aerated culture tubes such that they were in log phase with OD_600_>0.1 upon use the following day. Cells were then diluted to OD600=0.1. For experiments comparing genetic backgrounds, cells were grown for 3 h before measurement. For experiments testing the effects of drug stressors, stock solutions of the drugs were prepared in advance (10 mM bortezomib in ethanol, 10 mM hydrogen peroxide in water). The drugs were diluted to 50× concentrations, then added to cells 1:50 to reach 1× concentrations. The cells were grown for 5 h while shaking at 1,050 rpm before measurement. Fluorescence was measured on a BD Accuri C6 flow cytometer (BD Biosciences). All quantitative analysis was performed using MATLAB v8.6 (MathWorks). For the PSR, OSR, and *pRPN4:GFP* reporters, the GFP fluorescence measurements were normalized to forward scatter for each cell. For ERm-Deg and Cyto-Deg, GFP fluorescence measurements were normalized to RFP.

### lmmunoblotting

Samples for immunoblotting were collected at an OD_600_ of 0.4-0.8. One OD_600_ absorbance unit (1.25 - 2.5 ml) of cells were collected and lysed at 95°C in 15 µL 4× NuPage LDS Sample Buffer (Thermo Fisher Scientific) with 5% β-mercaptoethanol for 10 minutes. SDS-PAGE was performed on samples using 10% Tris-Glycine 1.5 mm gels. Samples were then transferred onto 0.45-µm nitrocellulose membranes (Bio-Rad) using a standard wet transfer protocol. Membranes were blocked with 5% milk in Tris-buffered saline with Tween for 1 h at room temperature. They were then stained with 1:2,000 Pierce monoclonal mouse anti-HA (26183; Thermo Fisher Scientific) primary antibody overnight at 4° C, followed by IRDye 800CW donkey anti-mouse (LiCor Biosciences) for 1 hat room temperature. Membranes were scanned using a LiCor Odyssey (LiCor Biosciences). Then, membranes were stained with 1:2,000 rabbit anti-hexokinase (H2035-01, US Biological) primary antibody for 3 h at room temperature, followed by IRDye 680RD goat anti-rabbit (LiCor Biosciences) for 1 hat room temperature. Membranes were then scanned again using a LiCor Odyssey. Scanned images were visualized with lmageJ (National Institutes of Health), and quantified with MATLAB v8.6 (MathWorks).

## Supplementary data files

**Supplementary data file 1: List of all sgRNA sequences and corresponding primers used in this study**. The sequences here are equivalent in name and sequence to those found in Alford et. al., 2021.

**Supplementary data file 2: Basal gene NNS’s and gene–stress interaction NNS’s for both stressors and both biological replicates used** in **this study**. Basal NNS values were computed by comparing the RNA and DNA counts of the barcodes for the sgRNAs of a given gene and averaging them. Gene-stress interaction NNS values were computed using the RNA counts of barcodes under stress conditions (either 20 µM bortezomib for 3 hr or 200 µM hydrogen peroxide for 30 min) compared to their RNA counts in unstressed conditions.

**Supplementary data file 3: Mean log**_**2**_ **fold change and standard deviation of the genes with the top and bottom 20 basal NNS’s**. Mean fold change was calculated by comparing the reporter RNA/DNA ratio under a given gene knockdown to the mean ratio. This value from each biological replicate was then averaged.

**Supplementary data file 4: Basal NNS’s for each sgRNA**. Basal NNS values were computed by comparing the RNA and DNA counts of the barcodes for each sgRNA and averaging them.

**Supplementary data file 5: sgRNA-stress NNS’s for both stressors used in this study**. NNS values were computed using the RNA counts of barcodes under stress conditions (either 20 µM bortezomib for 3 hr or 200 µM hydrogen peroxide for 30 min) compared to their RNA counts in unstressed conditions.

**Supplementary data file 6: Collection of files used to calculate gene NNS’s, GO category enrichment scores, and other pertinent information**. Raw data are contained in .mat files, and the .m functions can be used in MATLAB to generate scores. skiplist.m contains a list of genes that are omitted from the GO category analysis, and consists of tRNAs, retrotransposons, and mitochondrially encoded genes.

## Acknowledgements

We thank A. Raja Venkatesh, K. Le, L. Svoboda, and E. Tassoni-Tsuchida for helpful suggestions in preparation of the manuscript. This work was supported by the National Institute of General Medical Sciences of the US National Institutes of Health (grant No. T32GM007276 to J.J. Work, R01GM138689 to D. Pincus, and R01GM115968 to 0. Brandman).

## Competing interests

The authors declare no competing financial interests.

**Supplementary figure 1:**
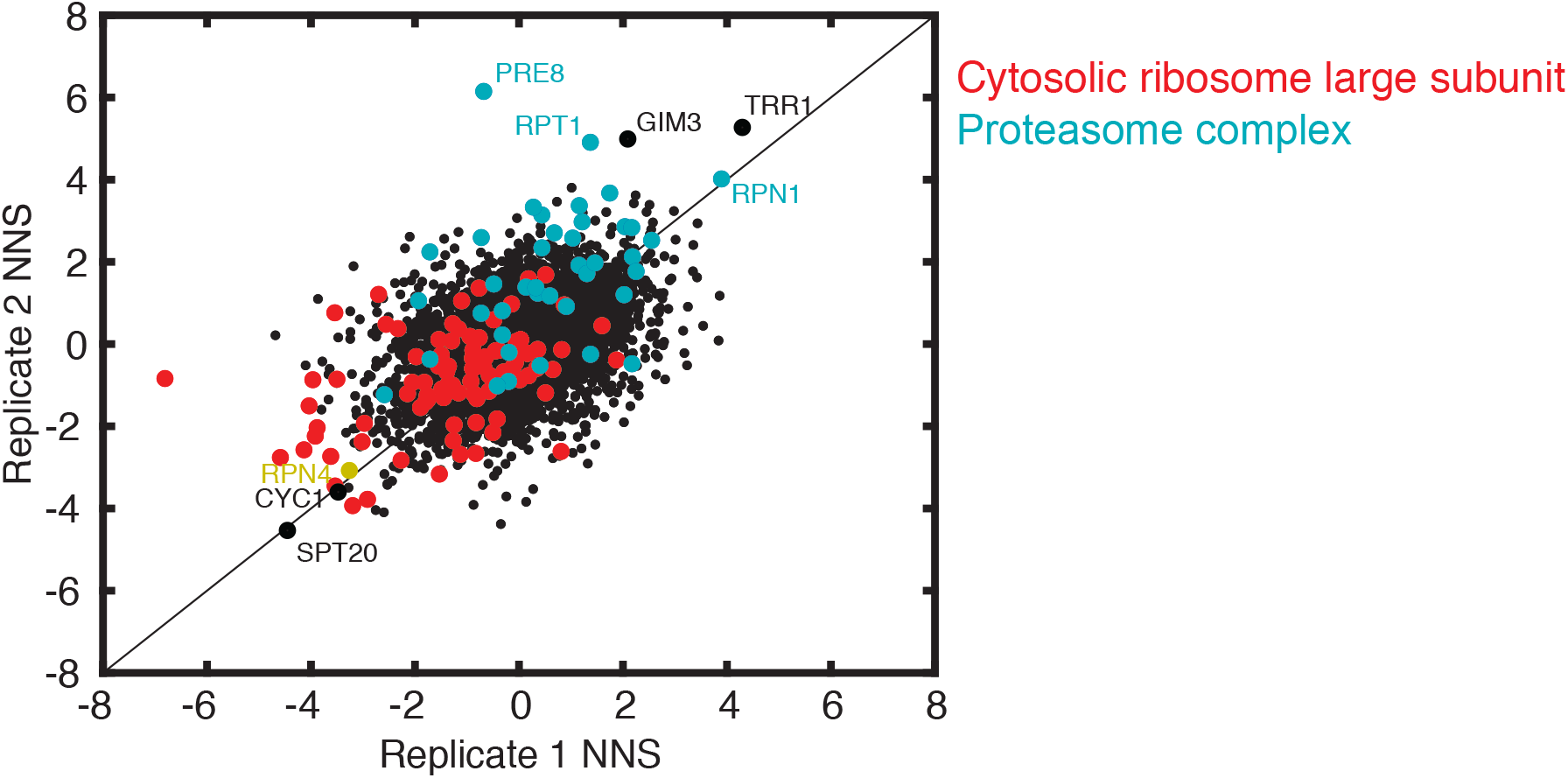
Biological replicates of ReporterSeq NNS values in basal conditions. *RPN4* and genes from the “cytosolic ribosome large subunit” and “proteasome complex” GO categories have been highlighted with color, and some genes of interest have been labeled.

**Supplementary figure 2:**
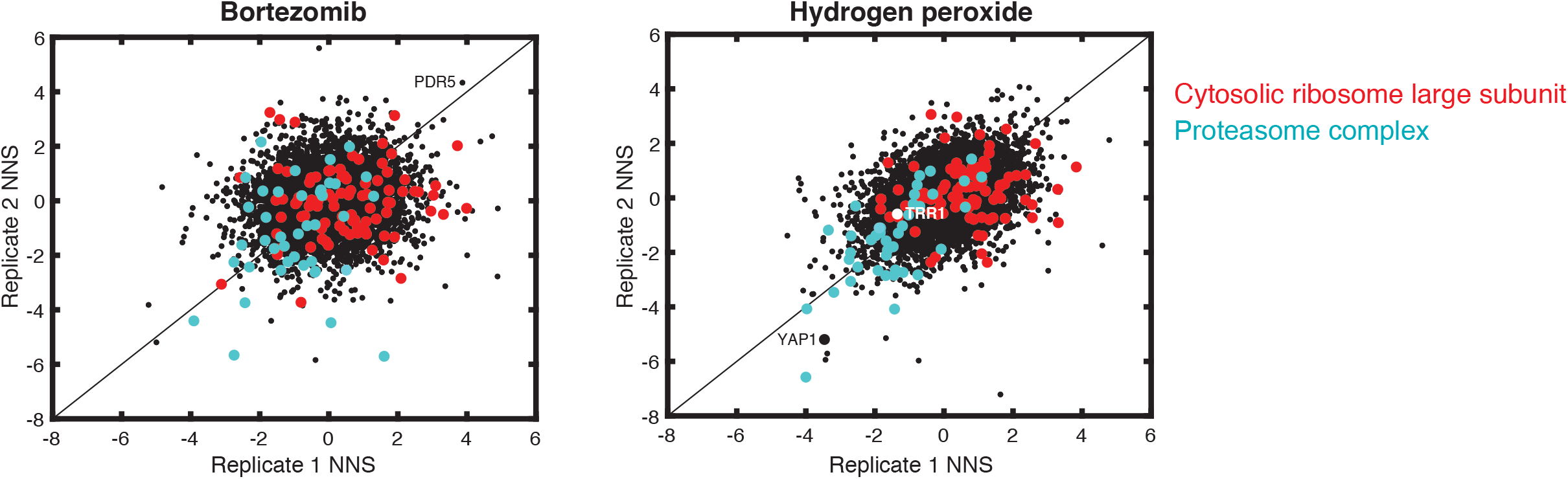
Biological replicates of ReporterSeq NNS values in stress conditions. Genes from the “cytosolic ribosome large subunit” and “proteasome complex” GO categories have been highlighted with color, and some genes of interest are labeled. Stress conditions were 3 hr 20 μM bortezomib (left) or 30 min 200 μM hydrogen peroxide treatment (right).

**Supplementary figure 4:**
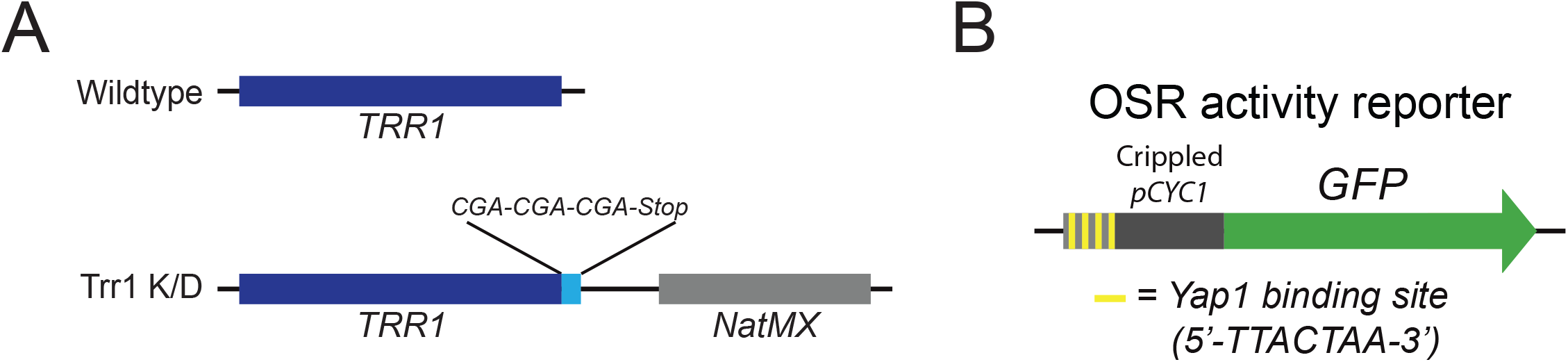
Knockdown strain and reporter diagrams. A: The Trr1 knockdown strain (Trr1 K/D) is created by inserting three CGA codons immediately upstream of the stop codon, with a NatMX cassette serving as a selectable marker. B: The OSR activity reporter is designed in near-identical fashion to the PSR activity reporter, but uses Yap1 binding sites instead of Rpn4 binding sites to report on the OSR instead of the PSR.

