## Supplementary Data File 6 for "ReporterSeq reveals genome-wide determinants of proteasome expression": getGeneScoresSG.m

```
function getGeneScores
%creates ReporterSeqScores, which includes
% basalScores (genes,replicates) array of basal genes scores
% basalFoldChange (genes,replicates) array of basal genes scores
% basalScoresMean average basal score per gene
% basalFoldChangeMean average basal fold change per gene
% basalSgRNAnum (gene-sgRNA) array of number of barcodes (sum of replicates)
% basalSgRNAScoreMean (gene-sgRNA) array of basal genes scores (per sgRNA) (average of replicates)
% basalSgRNAnames gene-sgRNA name pairs
% stressScores1 (gene,conditon) array scores, replicate one of stress condition
% stressScores2 (gene,conditon) array scores, replicate two of stress condition
% stressFoldChange1 (gene,conditon) array fold changes, replicate one of stress condition
% stressFoldChange2 (gene,conditon) array fold changes, replicate two of stress condition
% stressScoresMean (gene,condition) array of gene-stress interaction scores (average of replicates)
% stressFoldChangeMean (gene,condition) array of gene-stress interaction fold changes (average of replicates)
% stressSgRNAnum (gene-sgRNA,condition) array of number of barcodes (sum of replicates)
% stressSgRNAscore (gene-sgRNA,condition) array of gene-stress scores (per sgRNA) interactions (average of replicates)
% stressSgRNAnames (gene-sgRNA,condition) names
% L conditons
% g gene
% o orf

load readCounts;

basalRNADNAPairs=[];
for i=1:max(setNum)
    for DNAi=find(setNum'==i&strcmp(measurementType,'DNA')&strcmp(stressType,'none'))'
        for RNAi=find(setNum'==i&strcmp(measurementType,'RNA')&strcmp(stressType,'none'))'
            if (length(DNAi)>0&length(RNAi)>0)
                basalRNADNAPairs(end+1,:)=[RNAi DNAi];
            end
        end
    end
end
for i=1:rows(basalRNADNAPairs)
    basalScoresBC_(:,i)=getScore(D(:,basalRNADNAPairs(i,:)));
    basalFoldChangeBC_(:,i)=getFoldChange(D(:,basalRNADNAPairs(i,:)));
end

basalScoresBC=[nanmean(basalScoresBC_(:,1:4)')' nanmean(basalScoresBC_(:,5:8)')']; %average over technical replicates
basalFoldChangeBC=[nanmean(basalFoldChangeBC_(:,1:4)')' nanmean(basalFoldChangeBC_(:,5:8)')']; %average over technical replicates

%compute gene scores
[g,i]=unique(gene);
o=orf(i);
[basalSgScores,basalSgN,basalSgRNAnames]=barcodes2sgGeneScore(basalScoresBC,g,gene,sg);
[basalScores,basalN]=barcodes2geneScore(basalScoresBC,g,gene,1);
[basalFoldChange,basalFoldChangeN]=barcodes2geneScore(basalFoldChangeBC,g,gene,0);

%compute stress-gene interactions
for i=1:max(setNum)
    RNAbasal{i}=find(strcmp(stressType,'none')&setNum'==i&strcmp(measurementType,'RNA')); %pairs are technical replicates to be averaged
    RNAstress{i}=find(~strcmp(stressType,'none')&setNum'==i&strcmp(measurementType,'RNA'));
end

for i=1:length(RNAbasal)
    for basalRNAi=1:length(RNAbasal{i})
        for stressRNAi=1:length(RNAstress{i})
            stressScoresPerSetBC{i}(:,stressRNAi,basalRNAi)=getScore(D(:,[RNAstress{i}(stressRNAi) RNAbasal{i}(basalRNAi)]));
            stressFoldChangePerSetBC{i}(:,stressRNAi,basalRNAi)=getFoldChange(D(:,[RNAstress{i}(stressRNAi) RNAbasal{i}(basalRNAi)]));
        end
    end
end
%average technical replicates
for i=1:length(RNAbasal)
    if (size(stressScoresPerSetBC{i},3)>1)
        x=nan(size(stressScoresPerSetBC{i},1),size(stressScoresPerSetBC{i},2));
        x(:)=nanmean([LIN(stressScoresPerSetBC{i}(:,:,1)) LIN(stressScoresPerSetBC{i}(:,:,2))]');
        stressScoresPerSetBC{i}=x;
    else
        stressScoresPerSetBC{i}=stressScoresPerSetBC{i}(:,:,1);
    end
end
for i=1:length(RNAbasal)
    if (size(stressFoldChangePerSetBC{i},3)>1)
        x=nan(size(stressFoldChangePerSetBC{i},1),size(stressFoldChangePerSetBC{i},2));
        x(:)=nanmean([LIN(stressFoldChangePerSetBC{i}(:,:,1)) LIN(stressFoldChangePerSetBC{i}(:,:,2))]');
        stressFoldChangePerSetBC{i}=x;
    else
        stressFoldChangePerSetBC{i}=stressFoldChangePerSetBC{i}(:,:,1);
    end
end

%array scores into single table
c=0;
for i=1:length(stressScoresPerSetBC)
    for j=1:size(stressScoresPerSetBC{i},2)
        c=c+1;
        stressLabels{c}=stressType{RNAstress{i}(j)};
        stressScoresPerSetBCtable(:,c)=stressScoresPerSetBC{i}(:,j);
        stressFoldChangePerSetBCtable(:,c)=stressFoldChangePerSetBC{i}(:,j);
    end
end
%split scores by replicate
[stressLabels,i]=sort(stressLabels);
stressScoresPerSetBCtable=stressScoresPerSetBCtable(:,i);
stressScoresPerSetBCtable2(:,:,1)=stressScoresPerSetBCtable(:,1:2:end);
stressScoresPerSetBCtable2(:,:,2)=stressScoresPerSetBCtable(:,2:2:end);

stressFoldChangePerSetBCtable=stressFoldChangePerSetBCtable(:,i);
stressFoldChangePerSetBC2(:,:,1)=stressFoldChangePerSetBCtable(:,1:2:end);
stressFoldChangePerSetBC2(:,:,2)=stressFoldChangePerSetBCtable(:,2:2:end);


%compute gene scores
[stressScores1,stressN1]=barcodes2geneScore(stressScoresPerSetBCtable2(:,:,1),g,gene,1);
[stressScores2,stressN2]=barcodes2geneScore(stressScoresPerSetBCtable2(:,:,2),g,gene,1);
[stressS1sg,stressN1sg,stressGeneSgNames1]=barcodes2sgGeneScore(stressScoresPerSetBCtable2(:,:,1),g,gene,sg);
[stressS2sg,stressN2sg,stressGeneSgNames2]=barcodes2sgGeneScore(stressScoresPerSetBCtable2(:,:,2),g,gene,sg);

[stressFoldChange1,stressN1fc]=barcodes2geneScore(stressFoldChangePerSetBC2(:,:,1),g,gene,0);
[stressFoldChange2,stressN2fc]=barcodes2geneScore(stressFoldChangePerSetBC2(:,:,2),g,gene,0);


L=stressLabels(1:2:end);

%average over replicates
for i=1:length(L)
    x=nan(size(stressScores1(:,1)));
    stressScoresMean(:,i)=normalize(nanmean([stressScores1(:,i) stressScores2(:,i)]'));
    stressFoldChangeMean(:,i)=nanmean([stressFoldChange1(:,i) stressFoldChange2(:,i)]');
end

basalScoresMean=normalize(nanmean(basalScores'));
basalSgRNAScoreMean=normalize(nanmean(basalSgScores'));
basalSgRNAnum=sum(basalSgN');
stressSgRNAscore = (stressS1sg+stressS2sg)/2;

basalFoldChangeMean=(nanmean(basalFoldChange'));

stressSgRNAnum= (stressN1sg+stressN2sg);
stressSgRNAnames=stressGeneSgNames1;
save ReporterSeqScores basalScoresMean stressScoresMean g o L stressScores1 stressScores2 basalScores stressSgRNAnum stressSgRNAscore basalSgRNAnum basalSgRNAScoreMean stressSgRNAnames basalSgRNAnames  basalFoldChangeMean stressFoldChangeMean basalFoldChange stressFoldChange1 stressFoldChange2

end

function s=getScore(D)
min_cutoff = 10;  %if the sum of the raw counts is <min_cutoff, scores are assigned nan
sz_cutoff=10000;    %needs to be large enough to be a representative sample

SC = nan(size(D,1),2);
lD=log(1+D);

% compute scaling
scaling = median(maxk(lD(:,1),1000))/median(maxk(lD(:,2),1000));

ii=find(not(isnan(D(:,1)))& not(isnan(D(:,2))));
rd=round(5*(lD(ii,1)+scaling*lD(ii,2)))/5;  %this significantly speeds up computation of scores
v=unique(rd);
for i =1:max(size(v))
    j=0;
    ind=find(rd==v(i));
    if size(ind,1)>0
        while max(size(ind))<sz_cutoff
            j=j+1;
            ind=find(rd>=v(max(i-j,1)) & rd<=v(min(i+j,max(size(v)))));
        end
        sc = (lD(ii(ind),1)-scaling*lD(ii(ind),2))./(.0000000001+lD(ii(ind),1)+scaling*lD(ii(ind),2));
        ind=find(rd==v(i));
        SC(ii(ind),1) = (lD(ii(ind),1)-scaling*lD(ii(ind),2))./(.0000000001+lD(ii(ind),1)+scaling*lD(ii(ind),2)) - median(sc);
        temp=sort(sc,'descend');
        s1=std(temp);
        s2=std(temp(100:end-99));
        if s1>2*s2
            SC(ii(ind),2) = s1; %If this triggers, outliers are present
        else
            SC(ii(ind),2) = s2;
        end
    end
end
s=SC(:,1)./SC(:,2);
s(find(D(:,1)+D(:,2)<min_cutoff))=nan;

end

function [s,n,giList]=barcodes2sgGeneScore(x,ug,gene,sg)

ind=0;
gin=1;
for gi=1:length(ug)
    if (mod(gi,1000)==0)
        disp(gi)
    end
    ind=strcmpForSorted(gene,ind(end)+1);
    sgList=unique(sg(ind));
    d=x(ind,:);
    thisSg=sg(ind);
    for sgi=sgList
        giList{gin}=[ug{gi} ' sg' num2str(sgi)];
        s(gin,:)=nanmean(d(thisSg==sgi,:));
        n(gin,:)=sum(isfinite(d(thisSg==sgi,:)));
        gin=gin+1;
    end
end
s=normalizeMAT(s);
end

function [s,n]=barcodes2geneScore(x,ug,gene,doNormalization)
maxScore=2.5;
l=abs(LIN(x))>maxScore;
x(l)=maxScore.*sign(x(l));

ind=0;
for gi=1:length(ug)
    ind=strcmpForSorted(gene,ind(end)+1);
    c=0;
    if (mod(gi,1000)==0)
        disp(gi)
    end
    if (length(ind)>1)
        s(gi,:)=nanmean(x(ind,:));
    end
    n(gi,:)=sum(isfinite((x(ind,:))));
end
if doNormalization==1
    s=normalizeMAT(s);
end
end

function d=normalizeMAT(d)
for c=1:cols(d)
    d(:,c) = normalize(d(:,c));
end
end


function d=normalize(d)
x = d(isfinite(d));
d = d - mean(x);
x = d(isfinite(d));
d = d/std(x);
end

function r=strcmpForSorted(g,i)
thisG=g{i};
r=[];
while i<=length(g) & strcmp(g{i},thisG)
    thisG=g{i};
    r(end+1)=i;
    i=i+1;
end
end

function r=rows(d)
r=size(d,1);
end

function c=cols(d)
c=size(d,2);
end

function X=LIN(X)
X=X(:);
end


function s=getFoldChange(D)
l=min(D')>5;
D(~l,:)=nan;
D=log2(D);
s=D(:,1)-D(:,2);
s=s-nanmedian(s);
end
```
