## Supplementary Data File 6 for "ReporterSeq reveals genome-wide determinants of proteasome expression": getGOenrichment.m

```
function getGOenrichment
%creates ReporterSeqScoresGOenrichment.mat which contans
% v (genes,conditions) array of values
% vsat (genes,conditions) array of values with 0.5% saturated to prevent outliers from dominating ttest
% catN name of GO category
% catN2 longer form of GO category name that also includes index in catN
% catT type of GO category
% catG genes in GO category
% goCat GO cat ID number
% go2id hash that maps GO ID to index in catN
% c conditions
% g genes
% o orf
% tt (GO categories,conditons) array of log(1./pval)

d=load('ReporterSeqScores');

%remove mito chr genes, tRNAs, retrotransposons
skiplist=textread('skiplist.m','%s');
[ig,i]=intersect(upper(d.g),upper(skiplist));
l=true(size(d.g));
l(i)=false;
d.basalScoresMean=d.basalScoresMean(l);
d.basalScores=d.basalScores(l,:);
d.g=d.g(l);
d.o=d.o(l);
d.stressScoresMean=d.stressScoresMean(l,:);
d.stressScores1=d.stressScores1(l,:);
d.stressScores2=d.stressScores2(l,:);

g=d.g;
o=d.o;

load goNames
load goMap
goCats=keys(M);
v=[d.stressScoresMean d.basalScoresMean'];
vsat=saturateTopAndBottomMAT(v,0.5);
c=d.L;
c{end+1}='basal';

counter=0;
go2id = containers.Map;
for i=1:length(goCats)
    if (mod(i,100)==0)
        disp(i)
    end
    gList=M(goCats{i});
    [ig,ii]=intersect(g,gList);
    if isKey(goNames,goCats{i})
        counter=counter+1;
        temp=goNames(goCats{i});
        catN{counter}=temp{1};
        catT{counter}=temp{2};
        catG{counter}=gList;
        goCat{counter}=goCats{i};
        catN2{counter}=[temp{1} '| ' temp{2} '| ' goCats{i} '| ' num2str(counter)];
        go2id(goCats{i})=counter;
        for j=1:cols(v)
            ii=ii(isfinite(v(ii,j)));
            if (length(ii)>1)
                [ig,pval(counter,j)]=ttest2(vsat(:,j),vsat(ii,j));
                pV(counter,j)=mean(v(ii,j));
            else
                pval(counter,j)=nan;
            end
        end
    end
end
tt=log(1./pval);
tt(pV<0)=-tt(pV<0);
basalScores=d.basalScores;
save ReporterSeqScoresGOenrichment  v vsat catN catT catG goCat go2id catN2 c g o tt

end

function D=saturateTopAndBottomMAT(D,SaturatePercent)
for i=1:cols(D)
    D(:,i)=saturateTopAndBottom(D(:,i),SaturatePercent);
end
end

function D=saturateTopAndBottom(D,SaturatePercent)
if (sum(isfinite(D(:)))>100/SaturatePercent)
    S=D(isfinite(D));
    S=sort(S);
    Th=S(floor((1-SaturatePercent/100)*end));
    Tl=S(floor((SaturatePercent/100)*end));
    D(D>Th)=Th;
    D(D<Tl)=Tl;
end
end

function r=rows(d)
r=size(d,1);
end

function c=cols(d)
c=size(d,2);
end

function X=LIN(X)
X=X(:);
end
```
