## Supplementary Data File 6 for "ReporterSeq reveals genome-wide determinants of proteasome expression": makeFoldChangeTables.m

```
function makeFoldChangeTables
load('ReporterSeqScores.mat')

basalScoresMean(isnan(basalScoresMean))=0;
[ig,i]=sort(basalScoresMean);
l=[i(1:20) i(end-19:end)];

f=fopen('basalFoldChange.txt','w');
fprintf(f,'Gene\tmeanFoldChange(log2) \tstandardDeviationFoldChange\n');
for i=l
    fprintf(f,'%s\t%f\t%f\n',g{i},nanmean(basalFoldChange(i,:)),nanstd(basalFoldChange(i,:)));
end
fclose(f);

stressScoresMean(isnan(stressScoresMean))=0;
[ig,i]=sort(stressScoresMean);
l=unique([i(1:20,:) i(end-19:end,:)])';

f=fopen('stressFoldChange.txt','w');
fprintf(f,'Gene');
for i=1:length(L)
    fprintf(f,'\t%sreplicate1FoldChange(log2)\t%sreplicate2FoldChange(log2)',L{i},L{i});
end
fprintf(f,'\n');

for i=l
    fprintf(f,'%s',g{i});
    for j=1:size(stressFoldChange1,2)
        fprintf(f,'\t%f\t%f',stressFoldChange1(i,j),stressFoldChange2(i,j));
    end
    fprintf(f,'\n');
end
fclose(f);
```
