## Supplementary Data File 6 for "ReporterSeq reveals genome-wide determinants of proteasome expression": makeScoreTables.m

```
function makeSgRNAScoreTables
load('ReporterSeqScores.mat')

f=fopen('NNS.txt','w');
fprintf(f,'Gene\tbasalRep1\tbasalRep2\tH2O2Rep1\tH2O2Rep2\tbortezomibRep1\tbortezomibRep2\n');
for i=1:length(g)
    fprintf(f,'%s\t%f\t%f\t%f\t%f\t%f\t%f\n',g{i},basalScores(i,1),basalScores(i,2),stressScores1(i,1),stressScores2(i,1),stressScores1(i,2),stressScores2(i,2));
end
fclose(f);
```
