## Supplementary Data File 6 for "ReporterSeq reveals genome-wide determinants of proteasome expression": makeSgRNAScoreTables.m

```
function makeSgRNAScoreTables
load('ReporterSeqScores.mat')

f=fopen('basalGeneSgScores.txt','w');
fprintf(f,'Gene sgRNAnum\tnumBarcodes\taverageScore\n');
for i=1:length(basalSgRNAnames)
    fprintf(f,'%s\t%i\t%f\n',basalSgRNAnames{i},basalSgRNAnum(i),basalSgRNAScoreMean(i));
end
fclose(f);


f=fopen('stessGeneInteractionSgScores.txt','w');
fprintf(f,'Gene sgRNAnum');
for i=1:length(L)
    fprintf(f,'\t%sNumBarcodes\t%saverageScore',L{i},L{i});
end
fprintf(f,'\n');

for i=1:length(basalSgRNAnames)
    fprintf(f,'%s',basalSgRNAnames{i});
    for j=1:size(stressSgRNAnum,2)
        fprintf(f,'\t%i\t%f',stressSgRNAnum(i,j),stressSgRNAscore(i,j));
    end
    fprintf(f,'\n');
end
fclose(f);
```
