## Supplementary Data File 6 for "ReporterSeq reveals genome-wide determinants of proteasome expression": plotCategoryGenes.m

```
function x(s,c,stressi,cati)
gset=c.catG{cati};
[ig,i]=intersect(s.g,gset);
plotl(s.basalRIC,s.stressGeneInteraction(:,stressi),'.');
hold on
plotl(s.basalRIC(i),s.stressGeneInteraction(i,stressi),'.',s.g(i));
title(c.catN{cati})
xlabel('basal')
ylabel(s.L{stressi});
hold off
```
