## Supplementary Data File 6 for "ReporterSeq reveals genome-wide determinants of proteasome expression": skiplist.m

```
tP(UGG)Q
Q0010
ORI1
Q0017
15S_RRNA
tW(UCA)Q
Q0032
ORI8
COX1
AI1
AI2
AI3
AI4
AI5_ALPHA
AI5_BETA
ATP8
ATP6
ORI7
Q0092
ORI2
tE(UUC)Q
COB
BI2
BI3
BI4
ORI6
OLI1
tS(UGA)Q2
VAR1
Q0142
Q0143
Q0144
ORI3
ORI4
21S_RRNA
SCEI
tT(UGU)Q1
tC(GCA)Q
tH(GUG)Q
Q0182
tL(UAA)Q
tQ(UUG)Q
tK(UUU)Q
tR(UCU)Q1
tG(UCC)Q
tD(GUC)Q
tS(GCU)Q1
tR(ACG)Q2
tA(UGC)Q
tI(GAU)Q
tY(GUA)Q
tN(GUU)Q
tM(CAU)Q1
COX2
Q0255
tF(GAA)Q
tT(UAG)Q2
tV(UAC)Q
COX3
ORI5
tM(CAU)Q2
RPM1
Q0297
tA(AGC)D
tA(AGC)F
tA(AGC)G
tA(AGC)H
tA(AGC)J
tA(AGC)K1
tA(AGC)K2
tA(AGC)L
tA(AGC)M1
tA(AGC)M2
tA(AGC)P
tA(UGC)E
tA(UGC)G
tA(UGC)L
tA(UGC)O
tA(UGC)Q
tC(GCA)B
tC(GCA)G
tC(GCA)P1
tC(GCA)P2
tC(GCA)Q
tD(GUC)B
tD(GUC)D
tD(GUC)G1
tD(GUC)G2
tD(GUC)I1
tD(GUC)I2
tD(GUC)J1
tD(GUC)J2
tD(GUC)J3
tD(GUC)J4
tD(GUC)K
tD(GUC)L1
tD(GUC)L2
tD(GUC)M
tD(GUC)N
tD(GUC)O
tD(GUC)Q
tE(CUC)D
tE(CUC)I
tE(UUC)B
tE(UUC)C
tE(UUC)E1
tE(UUC)E2
tE(UUC)E3
tE(UUC)G2
tE(UUC)G3
tE(UUC)I
tE(UUC)J
tE(UUC)K
tE(UUC)L
tE(UUC)M
tE(UUC)P
tE(UUC)Q
tF(GAA)B
tF(GAA)D
tF(GAA)F
tF(GAA)G
tF(GAA)H1
tF(GAA)H2
tF(GAA)M
tF(GAA)N
tF(GAA)P1
tF(GAA)P2
tF(GAA)Q
tG(GCC)B
tG(GCC)D1
tG(GCC)D2
tG(GCC)E
tG(GCC)F2
tG(GCC)G1
tG(GCC)G2
tG(GCC)J1
tG(GCC)M
tG(GCC)O1
tG(GCC)P1
tG(GCC)P2
tG(UCC)Q
tH(GUG)E1
tH(GUG)E2
tH(GUG)G1
tH(GUG)G2
tH(GUG)H
tH(GUG)K
tH(GUG)M
tH(GUG)Q
tI(AAU)B
tI(AAU)D
tI(AAU)E1
tI(AAU)E2
tI(AAU)G
tI(AAU)I1
tI(AAU)I2
tI(AAU)L1
tI(AAU)L2
tI(AAU)N1
tI(AAU)N2
tI(AAU)P1
tI(AAU)P2
tI(GAU)Q
tI(UAU)D
tI(UAU)L
tK(CUU)C
tK(CUU)D1
tK(CUU)D2
tK(CUU)E1
tK(CUU)E2
tK(CUU)F
tK(CUU)G1
tK(CUU)G2
tK(CUU)G3
tK(CUU)I
tK(CUU)J
tK(CUU)K
tK(CUU)M
tK(CUU)P
tK(UUU)D
tK(UUU)G1
tK(UUU)G2
tK(UUU)K
tK(UUU)L
tK(UUU)O
tK(UUU)P
tK(UUU)Q
tL(CAA)D
tL(CAA)G1
tL(CAA)G3
tL(CAA)K
tL(CAA)L
tL(CAA)M
tL(CAA)N
tL(GAG)G
tL(UAA)B1
tL(UAA)B2
tL(UAA)D
tL(UAA)K
tL(UAA)L
tL(UAA)N
tL(UAA)Q
tL(UAG)J
tL(UAG)L1
tL(UAG)L2
tM(CAU)C
tM(CAU)Q1
tM(CAU)Q2
tN(GUU)C
tN(GUU)F
tN(GUU)G
tN(GUU)K
tN(GUU)L
tN(GUU)N1
tN(GUU)N2
tN(GUU)O1
tN(GUU)O2
tN(GUU)P
tN(GUU)Q
tP(UGG)L
tP(UGG)N1
tP(UGG)N2
tP(UGG)O1
tP(UGG)O3
tP(UGG)Q
tQ(UUG)B
tQ(UUG)C
tQ(UUG)D1
tQ(UUG)D2
tQ(UUG)D3
tQ(UUG)E1
tQ(UUG)E2
tQ(UUG)H
tQ(UUG)L
tQ(UUG)Q
tR(ACG)D
tR(ACG)E
tR(ACG)J
tR(ACG)K
tR(ACG)L
tR(ACG)O
tR(ACG)Q2
tR(UCU)B
tR(UCU)D
tR(UCU)E
tR(UCU)G1
tR(UCU)G2
tR(UCU)G3
tR(UCU)J1
tR(UCU)J2
tR(UCU)K
tR(UCU)M1
tR(UCU)M2
tR(UCU)Q1
tS(AGA)A
tS(AGA)B
tS(AGA)D1
tS(AGA)D2
tS(AGA)D3
tS(AGA)E
tS(AGA)G
tS(AGA)H
tS(AGA)J
tS(AGA)L
tS(AGA)M
tS(GCU)F
tS(GCU)O
tS(GCU)Q1
tS(UGA)Q2
tT(AGU)B
tT(AGU)C
tT(AGU)D
tT(AGU)H
tT(AGU)I1
tT(AGU)I2
tT(AGU)J
tT(AGU)N1
tT(AGU)N2
tT(AGU)O1
tT(AGU)O2
tT(UAG)Q2
tT(UGU)G1
tT(UGU)G2
tT(UGU)H
tT(UGU)P
tT(UGU)Q1
tV(AAC)E1
tV(AAC)E2
tV(AAC)G1
tV(AAC)G2
tV(AAC)G3
tV(AAC)H
tV(AAC)J
tV(AAC)K1
tV(AAC)K2
tV(AAC)L
tV(AAC)M1
tV(AAC)M2
tV(AAC)M3
tV(AAC)O
tV(CAC)D
tV(CAC)H
tV(UAC)B
tV(UAC)D
tV(UAC)Q
tW(CCA)G1
tW(CCA)G2
tW(CCA)J
tW(CCA)K
tW(CCA)M
tW(CCA)P
tW(UCA)Q
tX(XXX)D
tX(XXX)L
tY(GUA)Q
YAR009C
YAR010C
YBL005W-A
YBL005W-B
YBL100W-A
YBL100W-B
YBR012W-A
YBR012W-B
YCL019W
YCL020W
YDR034C-C
YDR034C-D
YDR098C-A
YDR098C-B
YDR170W-A
YDR210C-C
YDR210C-D
YDR210W-A
YDR210W-B
YDR261C-C
YDR261C-D
YDR261W-A
YDR261W-B
YDR316W-A
YDR316W-B
YDR365W-A
YDR365W-B
YER137C-A
YER138C
YER159C-A
YER160C
YFL002W-A
YFL002W-B
YGR027W-A
YGR027W-B
YGR038C-A
YGR038C-B
YGR109W-A
YGR109W-B
YGR161C-C
YGR161C-D
YGR161W-A
YGR161W-B
YHL009W-A
YHL009W-B
YHR214C-B
YHR214C-C
YIL082W
YIL082W-A
YJL113W
YJL114W
YJR026W
YJR027W
YJR028W
YJR029W
YLR035C-A
YLR157C-A
YLR157C-B
YLR227W-A
YLR227W-B
YLR256W-A
YLR410W-A
YLR410W-B
YML039W
YML040W
YML045W
YML045W-A
YMR045C
YMR046C
YMR050C
YMR051C
YNL054W-A
YNL284C-A
YNL284C-B
YOL103W-A
YOL103W-B
YOR142W-A
YOR142W-B
YOR192C-A
YOR192C-B
YOR343W-A
YOR343W-B
YPL060C-A
YPL257W-A
YPL257W-B
YPR137C-A
YPR137C-B
YPR158C-C
YPR158C-D
YPR158W-A
YPR158W-B
```
